# Plumbagin and oridonin reveal new CRM1 binding sites and NES-binding groove features

**DOI:** 10.1101/2020.08.05.237479

**Authors:** Yuqin Lei, Yuling Li, Yuping Tan, Da Jia, Qingxiang Sun

## Abstract

CRM1 is an important drug target in diseases such as cancer and viral infection. Plumbagin and oridonin, the herbal ingredients with known anti-cancer activities, were reported to inhibit CRM1-mediated nuclear export. However, their modes of CRM1 inhibition are unclear. Here, a multi-mutant of yeast CRM1 was engineered to enable the crystallization of these two small molecules in CRM1’s NES-binding groove. Each structure showed three inhibitor-binding sites, among which two are conserved in humans. Besides the known binding site, another site also participated in oridonin and plumbagin’s CRM1 inhibition. While the plumbagin-bound NES groove resembled the NES-bound groove state, the oridonin-bound groove revealed for the first time a more open NES groove, which may potentially improve cargo-loading through a capture-and-tighten mechanism. Our work thus provides a tool for CRM1 inhibitor crystallization, new insights of CRM1-cargo interaction, and a structural basis for further development of these or other CRM1 inhibitors.

## Introduction

In eukaryotic cells, the Chromosome Region Maintenance 1 protein (CRM1; exportin 1 or XPO1) is the major nuclear export receptor, which recognizes nuclear export signals (NES) – containing cargoes through a long and narrow surface groove called the NES-binding groove (or NES groove in short) [1, 2]. In order to export cargoes, CRM1 and cargo have to form a ternary complex with the small GTPase RanGTP in the nucleus, before traversing the nuclear pore complex to reach the cytoplasm [1–3]. Cytoplasmic factors RanGAP/RanBP1/RanBP2 disassemble the complex to release the cargo, meanwhile allowing CRM1/Ran to be recycled to nucleus for further rounds of nuclear export [3, 4].

Overexpression of CRM1 is observed in a variety of cancer cells and is often associated with poor clinical outcomes [5–7]. Due to CRM1’s function of nuclear export, several tumour suppressors and oncogenes such as p53, FOXO, and eIF4e, were abruptly exported to the cytoplasm in different cancer cells [8–10]. The outcome is often cancer-promoting and may cause resistance to anti-cancer therapies [11, 12]. Inhibiting CRM1 supress the majority of cancer hallmark features, CRM1 may thus be exploited as a broad-spectrum anti-cancer target [6]. Since nuclear exit of viral components are often essential for viral infections, CRM1 inhibitors may also be developed as broad-spectrum antiviral therapy [13].

Among the dozens of CRM1 inhibitors discovered, several were/are tested in clinical trials [5, 14]. The reported inhibitors were previously divided into four groups: bacterial products, herbal ingredients, fungal or animal inhibitors, and synthetic compounds [6]. All these inhibitors possess Michael acceptors that could covalently conjugate to a cysteine (C528) in the NES groove [15]. Prior structural studies illustrated how bacterial products (e.g. leptomycin B, Ratjadone) and synthetic NEIs (e.g. KPT-185, KPT-276, CBS9106, LFS-829) bind to CRM1, providing useful information for CRM1 biology and drug development [16–19].

Currently, little is known for the binding mode of plant inhibitors, such as plumbagin and oridonin [20, 21]. Plumbagin, which was derived from the plant *Plumbago scandens,* displayed anti-tumour activity against diverse cancer cells *in vitro* and *in vivo [22]*. It was reported that C528S mutant harbouring cells were resistant to growth inhibition by plumbagin [20]. The other anti-cancer natural product oridonin was derived from Chinese herbal medicine *Rabdosia rubescens* [23]. Nuclear accumulation of NPM1c+ and CRM1 was observed under oridonin treatment, though a direct interaction between CRM1 and Oridonin was not established [21]. We previously showed that both plumbagin and oridonin are direct inhibitors of CRM1 through pull down and cellular studies [24]. To characterize how the two plant inhibitors bind to CRM1, we carried out the following biochemical and biophysical studies.

## Results

### Crystal structure of plumbagin in complex with CRM1

To view the mode of plumbagin binding, we obtained its co-crystal structure using the previously reported Ran-RanBP1-CRM1 complex [4, 17]. This CRM1 construct contains *Saccharomyces cerevisiae* CRM1 residues 1-1055, mutation ^537^DLTVK^541^/GLCEQ, Y1022C, and 377-413 deletion, herein renamed as yCRM1a. In the crystal structure, two plumbagins are bound per CRM1 molecules, both covalently linked to a cysteine residue in the S configuration with well-defined densities (Figure 1A, B, S1). The C152-bound plumbagin lies in a shallow pocket formed by H4A, H4B, H5A, and H5B (Figure 1C). Except the conjugation, plumbagin forms only hydrophobic interactions with groove residues. There is a slight adjustment of CRM1 to accommodate the binding of plumbagin (Figure S2). In the C152-unliganded structure (4HAT), loop 5 (residues 204-207, between H5A and H5B) is packed towards H6. In the plumbagin-conjugated structure, loop 5 is flapped towards H4, forming hydrophobic interactions with this plumbagin (Figure S2). Another plumbagin which binds to C1022 is well buried in a channel formed by H19B, H20A, and H20B (Figure 1D). This plumbagin forms one hydrogen bond with the backbone of I963 on H19B and is intimately sandwiched between T1019 and the side chain of Y967, forming extensive hydrophobic interactions. In summary, the crystal structure shows that two plumbagins are covalently bound to C152 and C1022, respectively.

**Figure 1.**
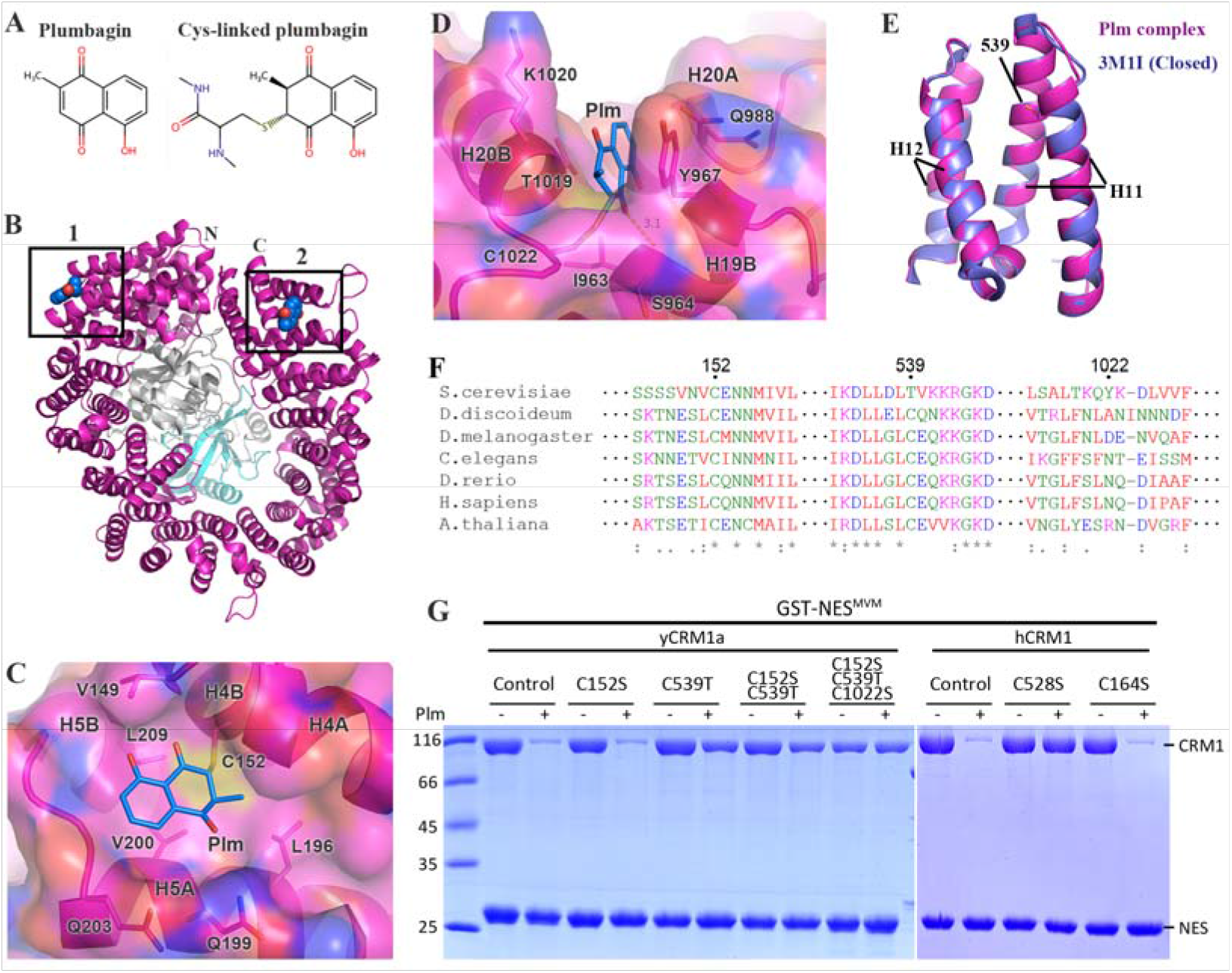
Crystal structure of plumbagin in complex with CRM1-Ran-RanBP1. A) Chemical structures of plumbagin (blue) and its cysteine-conjugated form. B) The overall structure of plumbagin (blue spheres) in complex with CRM1 (magenta), Ran (grey), and RanBP1 (cyan). C) C152-conjugated plumbagin (Plm, box 1 in 1B) in a pocket formed by H4 and H5. Residues within 4 Å distance are shown as sticks. D) C1022-conjugated plumbagin (box 2 in 1B) in a channel formed between H19B and H20. Residues within 4 Å distance are shown as sticks. E) Superimposition with the closed NES groove (3M1I, blue). The NES groove is highly similar to the closed groove (Cα RMSD 0.21 Å). C539 (in the Plm complex) and T539 (in 3M1I) are shown as sticks. The yeast CRM1 used (yCRM1a) contains a T539C mutation. F) Sequence alignment of CRM1 from different species. The numbers shown on top are the yeast numberings. yCRM1a contains a Y1022C mutation. G) Pull down of yCRM1a, hCRM1 or their mutants in the presence or absence of plumbagin. Human C528 corresponds to yeast C539. C164 in human corresponds to C152 in yeast. Plumbagin concentrations for yCRM1a and hCRM1 are 25 μM and 50 μM, respectively.

To our best knowledge, all known inhibitors of CRM1 are bound in the NES groove, forming covalent bonds with C539 (equivalent residue of human C528) and directly inhibiting NES binding. However, C539 is not bound with plumbagin in our crystal structure. Instead, the NES groove is highly similar to the groove-closed conformation (Figure 1E, 0.21 Å Cα RMSD to the 3M1I H11-H12 residues). Of the two covalently-linked cysteines, C152 but not C1022 is conserved (Figure 1F). In fact, C1022 is probably a cloning artefact since cysteine is not recorded in any known sequence of yeast (*S. cerevisiae*) or other species (Figure 1F), although C1022 existed in all our earlier works [16–18]. Through pull down, it was found that plumbagin’s inhibition of NES-binding was unchanged by C152S mutation, suggesting that C152 is not important for plumbagin’s NES inhibition (Figure 1G). A majority of inhibition was abolished by C539T mutation for yeast CRM1, suggesting that C539-conjugation plays a major role in plumbagin’s NES inhibition (Figure 1G). Compared with C152S/C539T, further mutation of C1022S completely abolished plumbagin’s NES inhibition, suggesting that C1022-conjugation also participates in NES inhibition, although the effect is minor (Figure 1G). For human CRM1 (hCRM1), which does not contain 1022 equivalent cysteine (being an N residue), C528S itself is sufficient to abolish plumbagin’s inhibition (even at a slightly increased plumbagin concentration), in agreement with the yCRM1a pull down results. (Figure 1G). In conclusion, the role played by C152, C539 and C1022 conjugations towards plumbagin’s NES inhibition were none, major, and minor, respectively.

### Structure-based design of CRM1 mutants to crystallize the inhibitors in the NES groove

The missing C539-bound plumbagin in our crystal structure might be due to RanBP1-enforced closure of CRM1’s NES groove (which became inaccessible to even excess of soaked plumbagin). Pre-incubating CRM1 with plumbagin, however, failed to yield any crystals. Two mutations were designed to open up the NES groove, namely yCRM1b (yCRM1a + Δ441-461) or yCRM1c (yCRM1a + Δ441-461, S553R, Q561E). These two

constructs showed reduced CRM1-Ran-RanBP1 complex affinity, and no protein crystals were obtained (Figure 2A, S3). Mutations by S27E based on yCRM1c (named yCRM1d) did increase the complex affinity (Figure 2B), however, remained non-crystallisable in the presence or absence of plumbagin, with or without re-screening. We suspected that crystal packing may be deteriorated after those mutations. Thus, two more residues (Q49E and A741T, based on yCRM1d) were further mutated, named as yCRM1e, to improve crystal packing (Figure 2C). All these mutations were designed to improve inter-molecular electrostatic interactions. For example, S27E and Q49E were designed to form electrostatic interaction with Ran in the same or adjacent asymmetric unit, respectively. A741T was designed to form hydrogen bonds with the symmetry-related CRM1 residue N305. The new mutant yCRM1e could be purified to a similar yield as yCRM1a and formed a good complex with Ran and RanBP1 (Figure S3).

**Figure 2.**
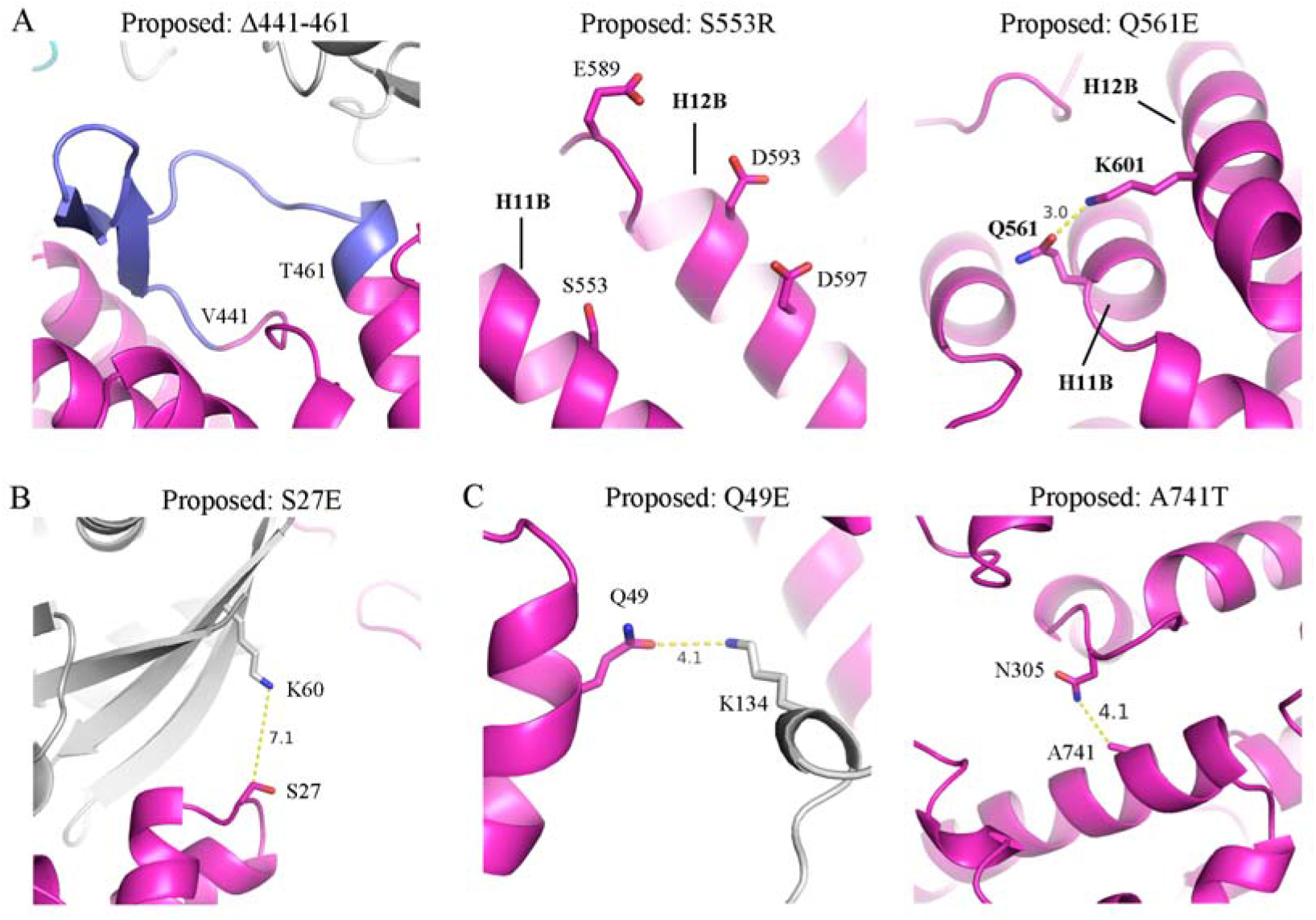
Mutations designed to enable C539-conjugation in co-crystals. A) Mutations designed to open-up the NES groove. CRM1, Ran, and RanBP1 are coloured magenta, grey, and cyan, respectively. The to-be-deleted H9 loop is coloured blue. Numbers represent the interatomic distances. In the absence of H9 loop, S553R and Q561E may facilitate the closure of H11B and H12B, restricting the open-close shuffling of this region. B) Mutation to increase the CRM1-Ran affinity through electrostatic interaction. C) Mutations to improve the crystal packing. These interactions are between molecules from adjacent symmetry mates.

### Crystal structures of plumbagin in complex with yCRM1e

With the help of yCRM1e, the crystal structure of C152/C539/C1022-conjugated plumbagin complex was successfully obtained with well-defined electron densities (Figure S4). Though the spacegroup is the same (and unit cell parameters are highly similar) (Table 1), the crystallization condition changed to a somewhat different condition (see methods) [17]. yCRM1e-C152 bound plumbagin displays subtle changes compared with the yCRM1a-C152 bound plumbagin, possibly due to higher B factors at this region (Figure 3A). The yCRM1e-C1022 bound plumbagin is highly similar to the yCRM1a-C1022 bound plumbagin conformation (Figure 3B). The third plumbagin indeed binds to the NES groove, mainly in the ◻3 pocket (Figure 3C). The base of this pocket is formed by residues A552 and I555, while L536, E540, K579, and F583 form the two sides of the groove. This plumbagin only forms hydrophobic interactions with CRM1. Its ketone/phenol pair is buried inside the pocket, forming intra-molecular hydrogen bonds. The lone ketone group is exposed to the solvent. In the ◻2 pocket and at a van der waals distance to the plumbagin molecule, a weak solitary electron density is observed and interpreted to be a DMSO molecule, which is the solvent used to dissolve plumbagin. As speculated, when H11A is aligned, plumbagin clashes heavily with the H12A in the yCRM1a complex (Figure 3D). The mutated residues largely formed the additional hydrogen bond or charge-charge interactions as we proposed (Figure S5). Acquired along multiple rounds of PCR reactions, an unintended mutation A51V is observed by electron density and verified by DNA sequencing. A51 or V51 is buried inside the protein and not involved in any inter-molecular contacts.

**Table 1.**
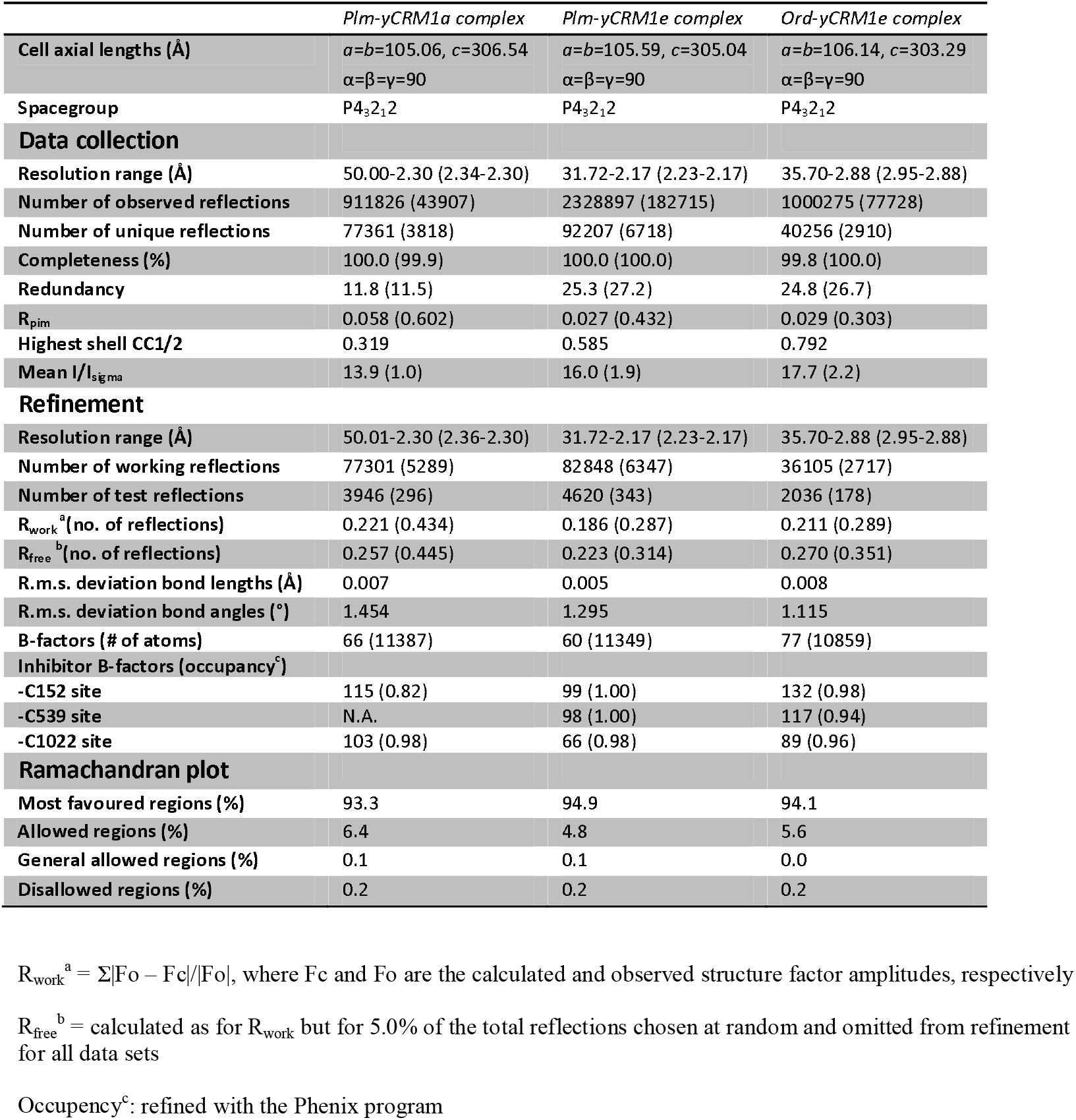
Data collection and structure refinement statistics.

**Figure 3.**
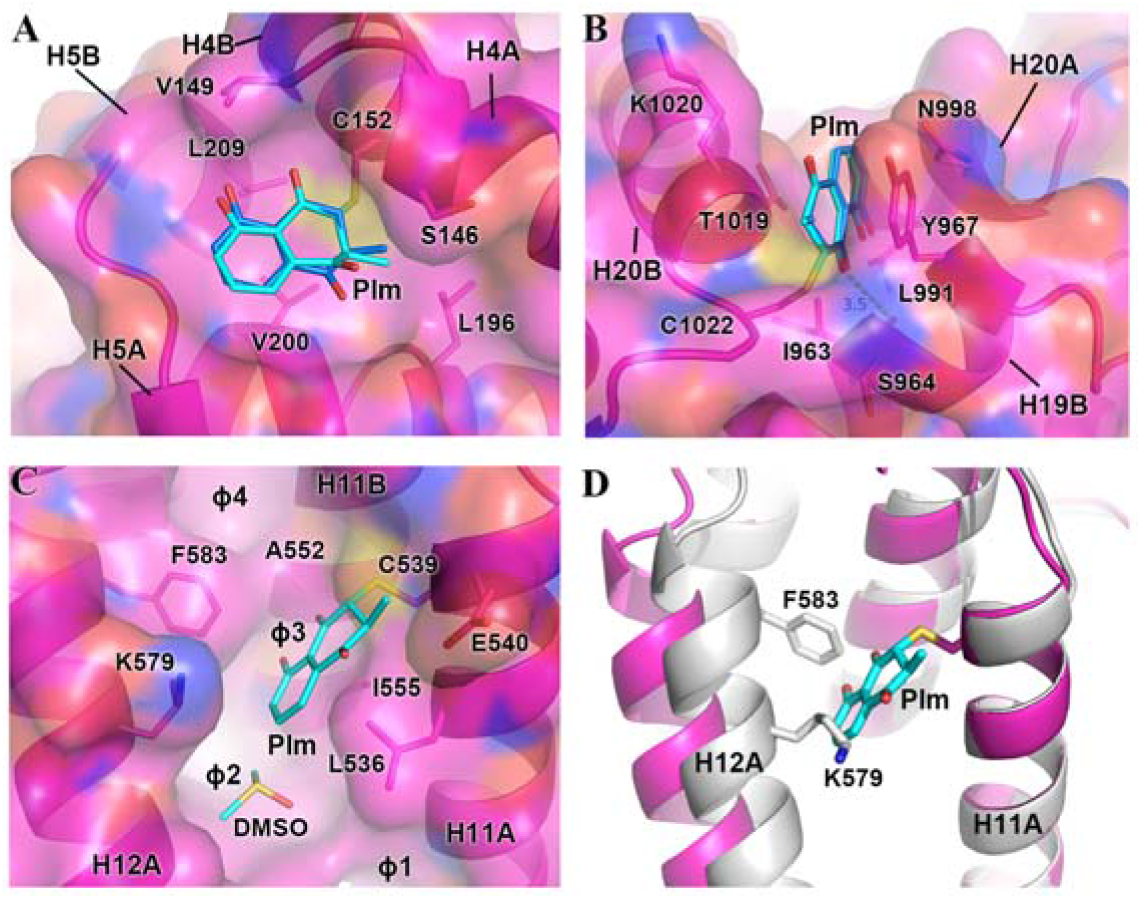
Crystal structures of yCRM1e in complex with plumbagin. A) C152-conjugated plumbagin (cyan) in the pocket formed by H4 and H5. Residues within 4 Å of plumbagin are shown as sticks. yCRM1a-bound plumbagin (blue) is superimposed for comparison. B) C1022-conjugated plumbagin in a channel formed between H19B and H20. Residues within 4 Å of plumbagin are shown as sticks. yCRM1a-bound plumbagin (blue) is superimposed for comparison. C) The NES groove with the bound plumbagin. Residues within 4 Å of plumbagin are shown as sticks. D) Alignment of yCRM1a (grey, no Plm-bound) with yCRM1e (magenta, Plm-bound) using the H11A residues (529-540). Plumbagin severely clashes with two residues on H12A in yCRM1a.

### Crystal structures of oridonin in complex with yCRM1e

Using the same strategy, the oridonin complex crystal structure was obtained, which also showed conjugations to C152, C539, and C1022 with well-defined density (Figure S6). C152-conjugated oridonin is orientated similarly to C152-conjugated plumbagin, surrounded by residues from H4A, H4B, H5A, and H5B (Figure 4A, S7). H4 loop and H5 loop are slightly different when bound to either plumbagin or oridonin. This oridonin additionally forms two hydrogen bonds with S146 and G204 in the pocket (Figure 4B). C1022-bound oridonin, however, binds very differently from C1022-bound plumbagin, mostly like because of being too bulky to fit into the narrow channel occupied by the plumbagin (Figure S8). Instead, this oridonin is more exposed to the solvent, forming three hydrogen bonds with the Y967 backbone and D968 side chain in the crystal structure (Figure 4C). C539-conjugated oridonin also binds to the ◻3 pocket in the NES groove, however, additionally occupies the ◻2 pocket due to being longer than plumbagin (Figure 4D). At this site, the residues that interact with oridonin are virtually the same as the residues interacting with plumbagin. Again, several ketone/hydroxyl groups in the oridonin are buried inside the NES groove, forming only intra-molecular hydrogen bonds. There is a void space between oridonin and the ◻2 pocket (Figure 4E), suggesting that this interaction is sub-optimal. Further extension of hydrophobic groups to fill the space may greatly enhance CRM1-binding affinity and selectivity. Similar to plumbagin, the binding to C539 and C1022, but not C152, participates in NES inhibition (Figure 4F). In agreement, the binding affinity towards MBP-NES^MVM^ (MVM NES is a supra-physiological NES that binds CRM1 in the absence of RanGTP [25]) was about 5 fold weaker for the C152/C539 double mutant compared with the C152/C539/C1022 triple mutant by microscale thermophoresis (MST) experiments (Figure S9).

**Figure 4.**
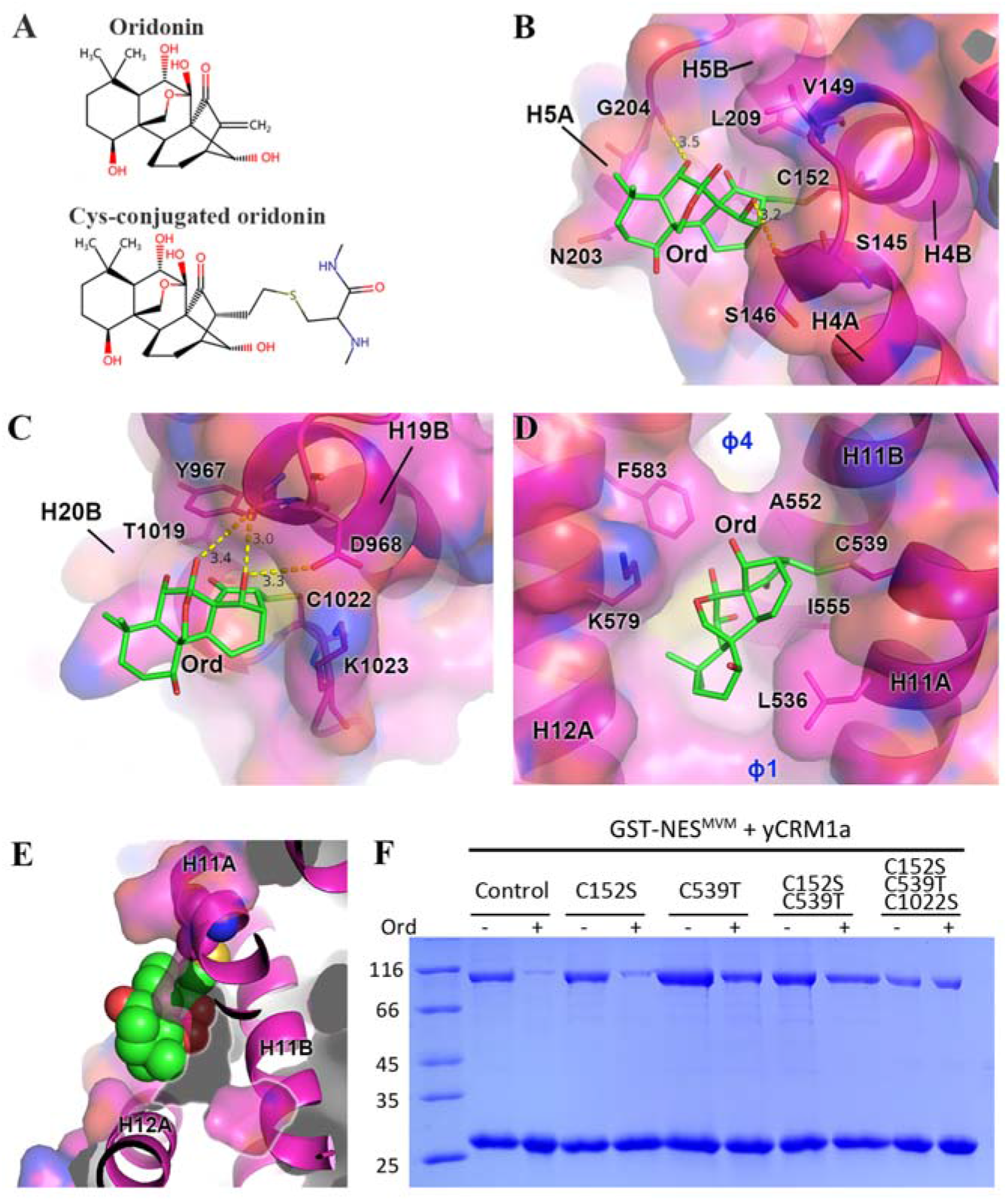
Crystal structures of yCRM1e in complex with oridonin. A) Chemical structures of oridonin and its cysteine-conjugated form. B) The C152 site with the bound oridonin (Ord). Hydrogen bonds are labelled as yellow dash lines. Residues within 4 Å of plumbagin are shown as sticks. C) The C1022 site with the bound oridonin. Residues within 4 Å of plumbagin are shown as sticks. D) The NES groove with the bound oridonin. Residues within 4 Å of plumbagin are shown as sticks. E) Illustration of the void space embedded by oridonin. Part of H11A (towards the reader) is clipped away in order to see the void space. Oridonin and C539 are shown as spheres. The void space is circled. F) Pull down of yCRM1a or its mutants in the presence or absence of oridonin. Oridonin concentration used is 60 μM.

### Comparison with the previously published structures

Next, the plumbagin- and oridonin-bound NES grooves were compared with several reported CRM1 structures, namely, the NES-bound structure (pdb:6CIT), the leptomycin-B (LMB)-bound structure (4HAT), the KTP-276-bound structure (4WVF), and the groove-closed structure (3M1I). The superimposition was performed by aligning the H12A residues (F572-E582) (Figure 5A). E540-E582 and V529-F572 Cα distances are used to illustrate the openness of the top or bottom part of the groove, respectively. E540 and V529 displacements based on the NES-bound structure are used to illustrate the H12A-relative movement of the top and bottom part of H11A, respectively. The plumbagin-bound groove closely resembles the NES-bound groove, displaying similar H11A orientation, least changes of E540-E582 and V529-F572 Cα distances, minor E540 and V529 displacement values, and negligible groove residue (529-582) Cα RMSDs (0.2Å) (Figure 5B, C). In contrast, the oridonin-bound groove conformation is different from all the structures shown. The top part of the groove is 2.5 Å wider than the NES-bound groove, which is previously thought to be the most widely open groove state (Figure 5B). The scale of groove-widening is substantial, considering that the NES-bound groove is only about 4 Å wider than the closed groove. The groove-widening at the top is necessary to accommodate this oridonin (Figure S10). Though the openness of the groove bottom is the same as that of the NES-bound groove, the V529 position is changed by 1.1Å along the helical direction of H11A, suggesting a H12A-centered circular motion. In summary, the plumbagin-bound groove resembles the NES-bound groove, while the oridonin-bound groove is even wider than the NES-bound groove.

**Figure 5.**
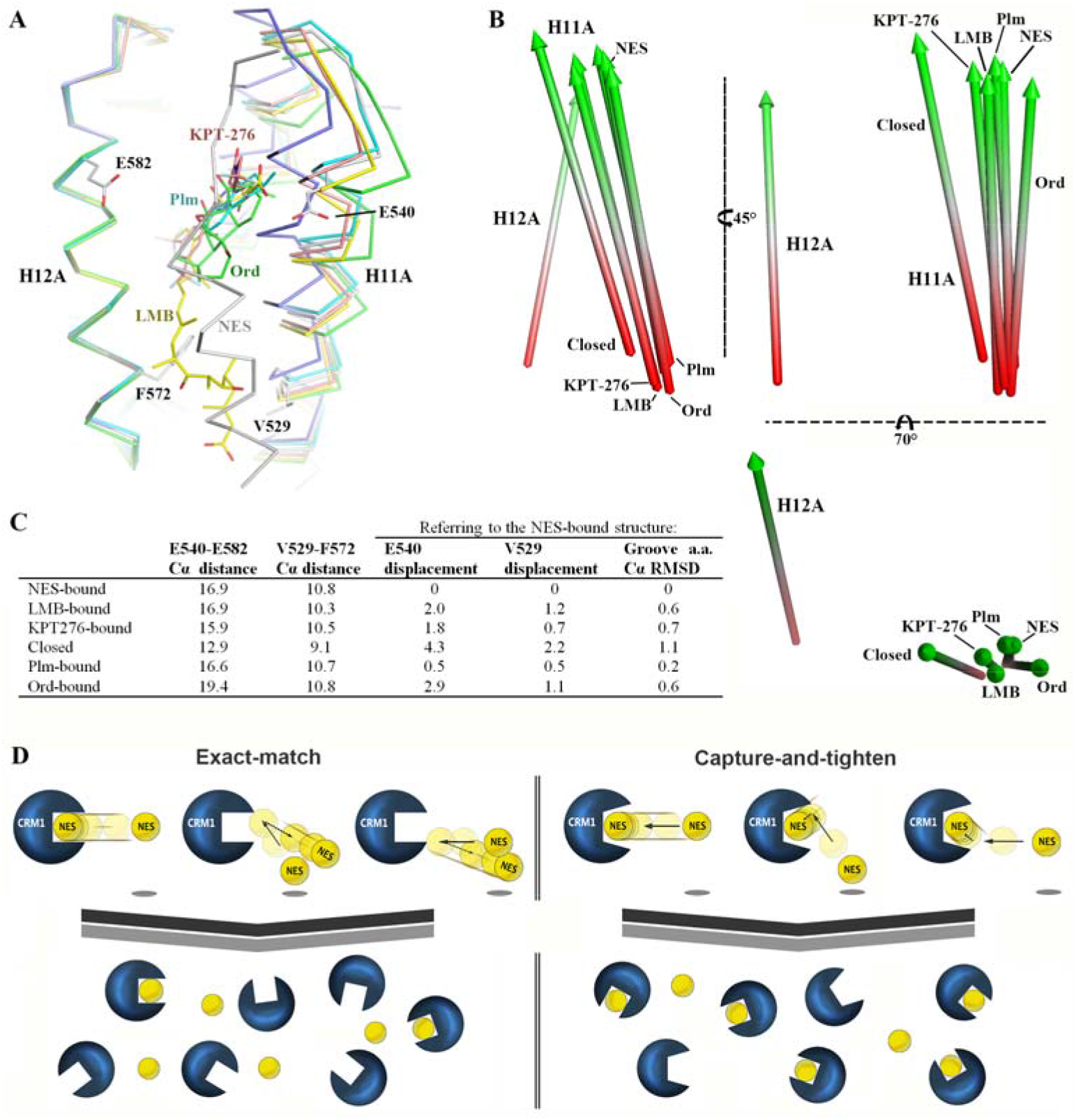
An observation of greater groove dynamics and a potential mechanism of cargo-loading. A) Superimposition of several CRM1 crystal structures including the NES-bound (grey), the LMB-bound (yellow), the KPT276-bound (brown), the groove-closed (blue), the plumbagin-bound (cyan), and the oridonin-bound (green) structures. All the structures in this figure are aligned by H12A residues F572-E582. V529, E540, F572, and E582 are shown as sticks. B) Helix orientations of H11A and H12A are simplified by spears pointing towards the carboxyl end of each helix. Red colour represents the amino end of each helix. C) Distances (Å) or RMSD values (Å) of the structures shown in panel A. The E540 (or V529) displacement values are the distance differences between E540 (or V529) of a specific structure and that of the NES-bound structure. Groove a.a. Cα RMSD stands for the root mean square deviation of groove residue (529-582) Cαs between a specific structure and the NES-bound structure. D) Comparison between two mechanisms of cargo-loading in the nucleus. Ran (probably also RanBP3) is bound to CRM1 but omitted for simplicity. On the left, the exact-match mechanism only accepts incoming cargoes from the straight-in direction. For the capture-and-tighten mechanism on the right, cargoes from other directions or arriving slightly off-cantered could also be captured through weak but adhesive interactions. The weak complexes could then undergo an interaction optimization step to form tight complexes. Compared with the exact-match mechanism, having a more open NES-groove could reduce the rejection rate of incoming cargoes and increase the efficiency of complex formation.

Previously, the H11A and H12A helices were thought to open and close in a 2-dimensional space, resulting in three states including the closed (3M1I), the half-open (KPT-276-bound) or the open (NES-bound) state [16, 18]. Here it is discovered that the groove dynamics are much more complicated than expected. The oridonin-bound and groove-closed structures are the two extremes of the groove conformations, while the other grooves (inhibitor- or NES-bound) could be considered as slightly deviating intermediates. Structurally morphing the oridonin-bound and groove-closed structures showed a greater flexibility (6.5 Å Cα distance difference between E540-E582) at the top part of the groove compared to the bottom part of the groove (1.7 Å Cα distance difference between V529-F572) (Movie 1). In addition to the previously known open-close motion in the plane defined by H11A and H12A, there is also an H12A-parallel motion in the plane and an H12A-orthogonal motion perpendicular to the plane (Movie 1). Thus, the NES groove dynamics could be redefined as ‘complex 3-dimonsional rearrangements featuring greater flexibility at the top (C528 end) of the groove’.

## Discussion

Plumbagin and oridonin are natural products that possess validated anti-cancer activities [23, 26]. They were reported to bind to several other cellular targets, such as ThyX[27], NLRP3[28], STAT3[29], AKT[30], Nucleolin[31], and AML1-ETO[32], yet no co-crystal structures have been reported to our best knowledge. Though LMB and SINE compounds (Small Inhibitors of Nuclear Export, including KPT-185, KPT-276, KPT-330, etc.) could be easily crystallized in complex with CRM1, the same system was incompatible for obtaining the crystal structures of plumbagin and oridonin. We suspected that the groove conformation was not suitable for the binding of these two molecules in that particular complex. Simple mutations to open up the NES groove meanwhile maintaining the complex affinity did not produce protein crystals. This was possibly due to the increased crystal packing energy, as the new complex with inhibitors might be slightly re-shaped and become less packable. Indeed, further mutations to improve crystal packing readily produced C539-conjugated crystals of plumbagin and oridonin in the same spacegroup but a different crystallization condition. Our experience is that the engineered mutant was more likely (about 80% crystals diffracted to 3.0 Å) to diffract to a high resolution compared with the original proteins (about 20% crystals diffracted to 3.0 Å). Thus, this structure-guided design of packing optimization could be potentially applied in other projects to enhance crystallizability and/or crystal resolution. In our opinion, mutagenesis to improve charge-charge interaction (even long-range) is more preferred than to improve hydrophobic interactions, because the latter may undesirably disrupt interaction due to steric clashes.

This is the first time observing CRM1 binding pockets other than the NES groove. It should be noted that the C152 and C1022 sites are not the FG-binding pockets reported earlier [33, 34]. Though C1022 does not exist in nature, its modification by plumbagin/oridonin was shown to inhibit NES binding slightly through pull down and MST. This site is far from the NES-binding site, but is very close to the C-terminus (Figure S11). When bound to NES and RanGTP, CRM1’s N-terminal domain and C-terminus contact each other, resulting in a more compact, ring-like CRM1 architecture. This N-C interaction is important for NES-groove opening by simulation and NES-binding by pull-down [35, 36]. The C1022 conjugation might potentially disrupt the N-C interaction, explaining the observed partial NES inhibition. The fact that C152 was able to bind to both plumbagin and oridonin, suggests that it may be a binding hotspot for more natural products or even metabolites in cells. Though C152 is not critical for NES-binding, modification of this site *in vivo* might also have biological consequences and may warrant further studies.

Interestingly, not all the surface-exposed cysteine residues are covalently conjugated by the two compounds. There are a few surface-exposed cysteine residues, namely, C152 (exposed surface area 8.1 Å^2^ by PISA server), C539 (11.3-12.6 Å^2^ depending on the groove state), C840 (27.6 Å^2^), C890 (7.9 Å^2^), C1022 (7.0 Å^2^). Though C840 and closed-groove C539 are more surface-exposed, they are not bound by inhibitors in the crystal structures. For C840 and C890, there are a greater number of acidic residues in the vicinity, which might inhibit the conjugation reaction [37]. Another reason might be that the surrounding environment clashes with the incoming inhibitors, as was explained above for C539 when the NES groove is closed. Besides, it is also possible that the C840 or C890-conjugated inhibitors do not form sufficient contacts with the surrounding pocket, hence de-conjugation occurred at a high speed (Michael addition is reversible [37]). The last two reasons may also explain why C152 or C1022 was not conjugated with all other CRM1 inhibitors.

Our crystal structures show that both plumbagin- and oridonin-bound grooves are different from the previously characterized inhibitor-bound grooves. While the plumbagin-bound groove is highly similar to the NES-bound groove, the oridonin-bound groove is even wider than the NES-bound groove, especially at the top of the groove. These observations explain why it is necessary for the engineered mutations in order to crystallize the two inhibitors in the groove. The oridonin-bound structure further expanded our knowledge of the NES groove dynamics. The groove simulation based on the oridonin-bound and groove-closed conformations shows a more complicated groove motion, rather than a simple open-close 2D motion. The top and bottom parts of the groove are of different flexibility. In addition, the H11A could also move in two other previously uncharacterized directions. It should be noted that H11B (and to a decreasing extent for H10, H9 …) also changes concomitantly while H11A moves, possibly impacting the architecture and rigidity of CRM1 (Movie 1). The groove information presented here should be considered for further structure-guided drug development.

Since wide-type human CRM1 could be inhibited by oridonin in vitro and in cells [21, 24], its NES-groove must be able to open wide-enough to accommodate oridonin binding. Thus, the observed wider NES-groove should exist in nature (not only in the crystal structure) and probably has a biological significance. Our hypothesis is that it may increase the efficiency of cargo loading. In a binding event, if CRM1’s NES groove could maximally open as big as the size of an NES, only the incoming cargoes from the straight-in direction would be captured (Figure 5D). Cargoes from other directions or arriving at the edge of the groove might be rejected before forming a tight complex. On the other hand, having a more open NES groove helps establishing weak but adhesive initial contacts between CRM1 and the cargo, even though the incoming cargoes were not in the straight-in direction. A further interaction-optimization step would then form a tight complex. Thus, having a more open NES-groove could reduce the rejection rate of incoming cargoes and improve the success rate of complex formation. Take an analogy, catches a flying baseball with the hand opening only as big as the ball size is probably not a good idea. Similarly, to pick up marbles with a pair of chopsticks, the initial openness of the pick-up-end should be slightly larger than the size of the marble. Thus, regardless of fast-moving (catching a ball) or slow-moving (picking up marbles), the ‘exact-match’ mechanism is always less efficient than the ‘capture-and-tighten’ mechanism.

The performance of both mechanisms is probably similar if the interaction surface is rather flat. For CRM1-cargo interaction (average groove depth about 8 Å), and probably other binary interactions involving deep penetrations, the later mechanism may be potentially more advantageous and more employed in biological systems.

## Materials and Methods

### Chemicals

Oridonin and plumbagin were purchased from Selleck (Shanghai, China) and initially dissolved at 100 mM concentration in DMSO.

### Cloning, expression and purification of proteins

The yeast CRM1 and their mutants were cloned into a PGEX-4t-1 expression vector, with a GST tag and incorporating a TEV cleavage site. CRM1 was expressed in E. coli BL21 (DE3), and grow in TB medium. CRM1 was induced by 500 μM IPTG at 18 °C overnight. Cells were harvested and re-suspended in lysis buffer (50 mM Tris pH 7.5, 200 mM NaCl, 10% glycerol, 2 mM DTT and 1 mM PMSF). The protein was loaded onto a GST column and was incubated with TEV (1mg:50mg) at 4 °C overnight to remove the GST tag. Next, the digested protein was eluted by a buffer containing 50 mM Tris pH 7.5, 200 mM NaCl, 2 mM DTT and 10% glycerol. The elution was purified by a Superdex 200 gel filtration column on the ÄktaPure (GE Healthcare) using the gel filtration buffer (20 mM Tris pH 7.5, 200 mM NaCl, 10% glycerol, 2 mM DTT). The protocol for expression and purification of human Ran (hRan) and yeast RanBP1(yRanBP1) was described previously [38]. Ran^L182A/Q69L^ (GTP-charged) was used for crystallization experiment [39].

### Crystallization of CMR1-Ran-RanBP1 complex and structure determination

The complex was prepared by mixing CRM1, Ran and RanBP1 at 1:3:2 molar ratio, and purified by Superdex 200 using a buffer containing 10 mM Tris pH 7.5, 100 mM NaCl, 5 mM MgCl_2_, and 1 mM EGTA. The purified complex was collected and concentrated to 5.5 mg/ml. The yCRM1a complex is crystallized as previously reported [17]. The yCRM1e complex was not crystallisable in the previous condition (18% PEG 3350, 200mM Ammonium Nitrate, 100mM Bis-Tris pH 6.6), but in the crystallization solution containing 0.12 M Monosaccharides (20 mM D-Glucose; 20 mM D-Mannose; 20 mM D-Galactose; 20 mM L-Fructose; 20 mM D-Xylose; 20 mM N-Acetyl-D-Glucosamine), 0.1 M buffer system 1 pH 6.5 (sodium HEPES and MOPS), and 50 % Precipitant Mix 2 (40% v/v Ethylene glycol; 20 % w/v PEG 8000). The crystals were soaked separately with oridonin or plumbagin for 2 h at 16 °C. Before freezing, the crystals were briefly dipped into the cryo-protectant containing 0.12 M Monosaccharides, 0.1 M buffer system 1 pH 6.5, 50 % Precipitant Mix 2, 10% glycerol (v/v), and 2.5 mM inhibitor.

X-ray diffraction data was collected at Shanghai Synchrotron Radiation Facility (SSRF) beamline BL17U1 and BL19U1 [40]. Coordinates of yCRM1-hRan-yRanBP1 (pdb code: 4HAT) were used as the search model using Molrep program [41], and refined using the program Refmac5 with Translation / Libration / Screw (TLS) refinement [42]. The data collection and refinement statistics were provided in Table 1.

### GST-Pull down assay

GST-NES^MVM^ or GST-yRanBP1 was immobilized on GSH beads. CRM1 proteins were incubated with different concentrations of compounds for 1 h at 25 °C. Different soluble proteins were incubated with the immobilized proteins with rotation in a total volume of 500 μL for 1 h at 4 °C. After three washing steps, bound proteins were separated by SDS/PAGE and visualized by Coomassie Blue staining. Each experiment was repeated at least twice and checked for consistency. The pull down buffer contains 20 mM Tris pH 8.0, 200 mM NaCl, 10% glycerol, 2 mM MgCl_2_, 0.005% Triton-X100.

## Data availability

The coordinates and structure factors were deposited in pdb with the accession codes : 5YSU, 6M60 and 6M6X.

## Acknowledgments

We thank funding from 1.3.5 project for disciplines of excellence, West China Hospital, Sichuan University.

## Conflict of interest

The authors declare no conflict of interest.

## Notes

### Competing Interest Statement

The authors have declared no competing interest.

