## Supplemental file for "Plumbagin and oridonin reveal new CRM1 binding sites and NES-binding groove features"

Running title: Crystal structures of oridonin and plumbagin in complex with CRM1

**This file includes figure S1-S11.**

**Supplemental Figures**

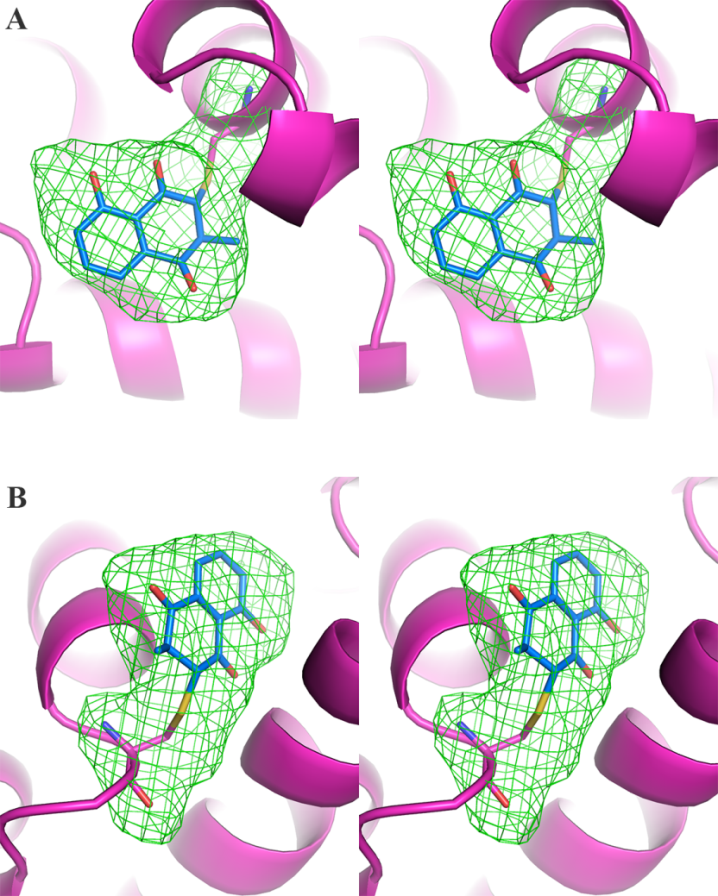

Figure S1. Stereo view of the Fo-Fc SA omit maps (green mesh) contoured at 4σ level. A) omit map for plumbagin and the conjugated cysteine C152. CRM1 is shown as magenta cartoon representation. B) omit map for plumbagin and the conjugated cysteine C1022.

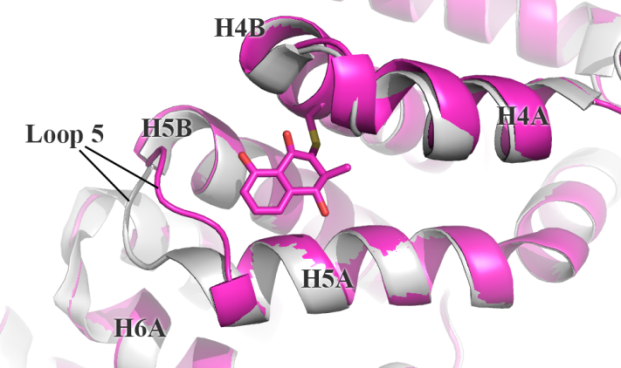

Figure S2. Comparison of C152-unliganded (4HAT, grey) and plumbagin-conjugated (magenta) CRM1 structures. Loop 5 is swung from H6 to H4 upon plumbagin binding.

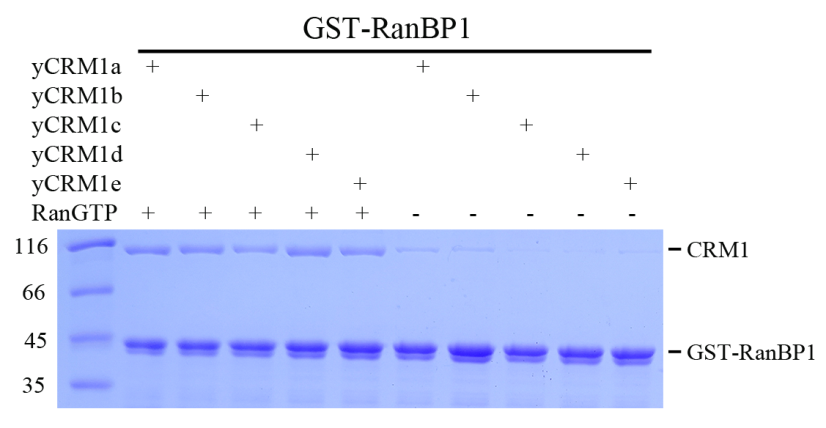

Figure S3. GST-RanBP1 pull down of different CRM1 (1 µM) constructs in the presence or absence of RanGTP (1 µM).

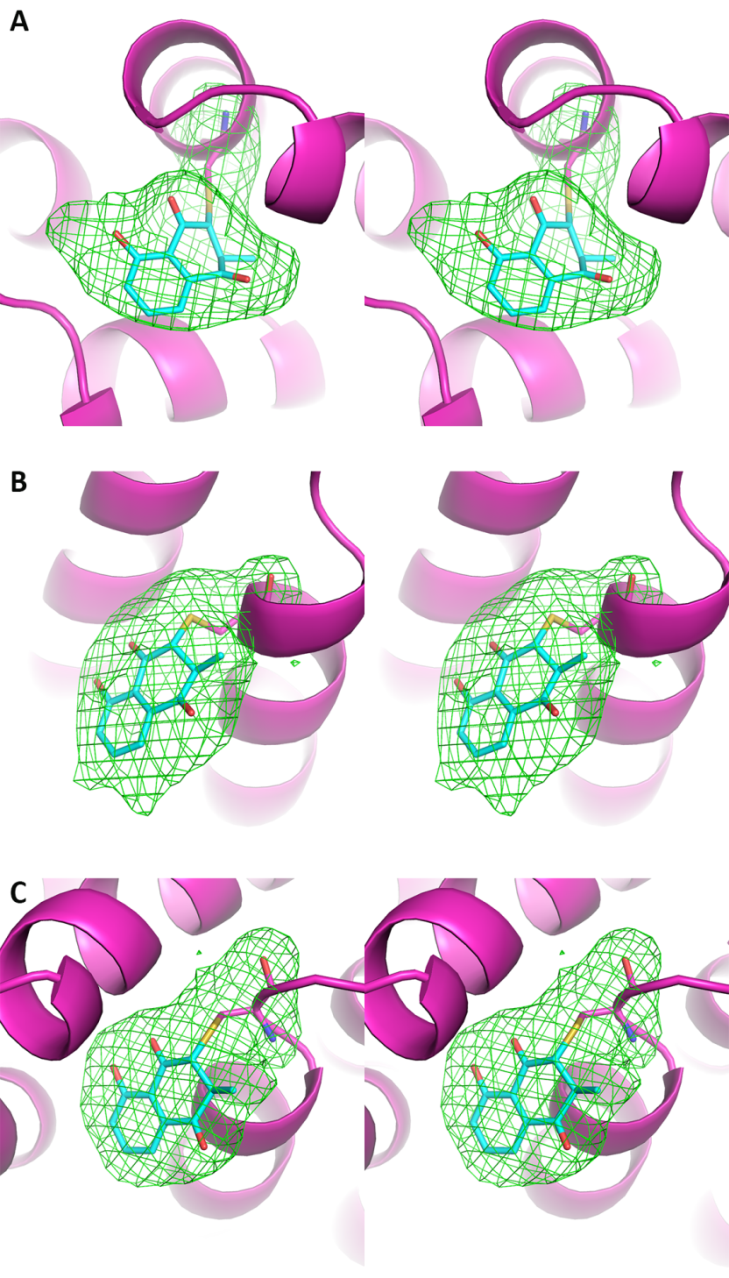

Figure S4. Stereo view of the Fo-Fc SA omit maps (green mesh) contoured at 4σ level. A) omit map for plumbagin and the conjugated cysteine C152. CRM1 is shown as magenta cartoon representation. Plumbagin is shown as cyan sticks. B) omit map for plumbagin and the conjugated C539. C) omit map for plumbagin and the conjugated C1022.

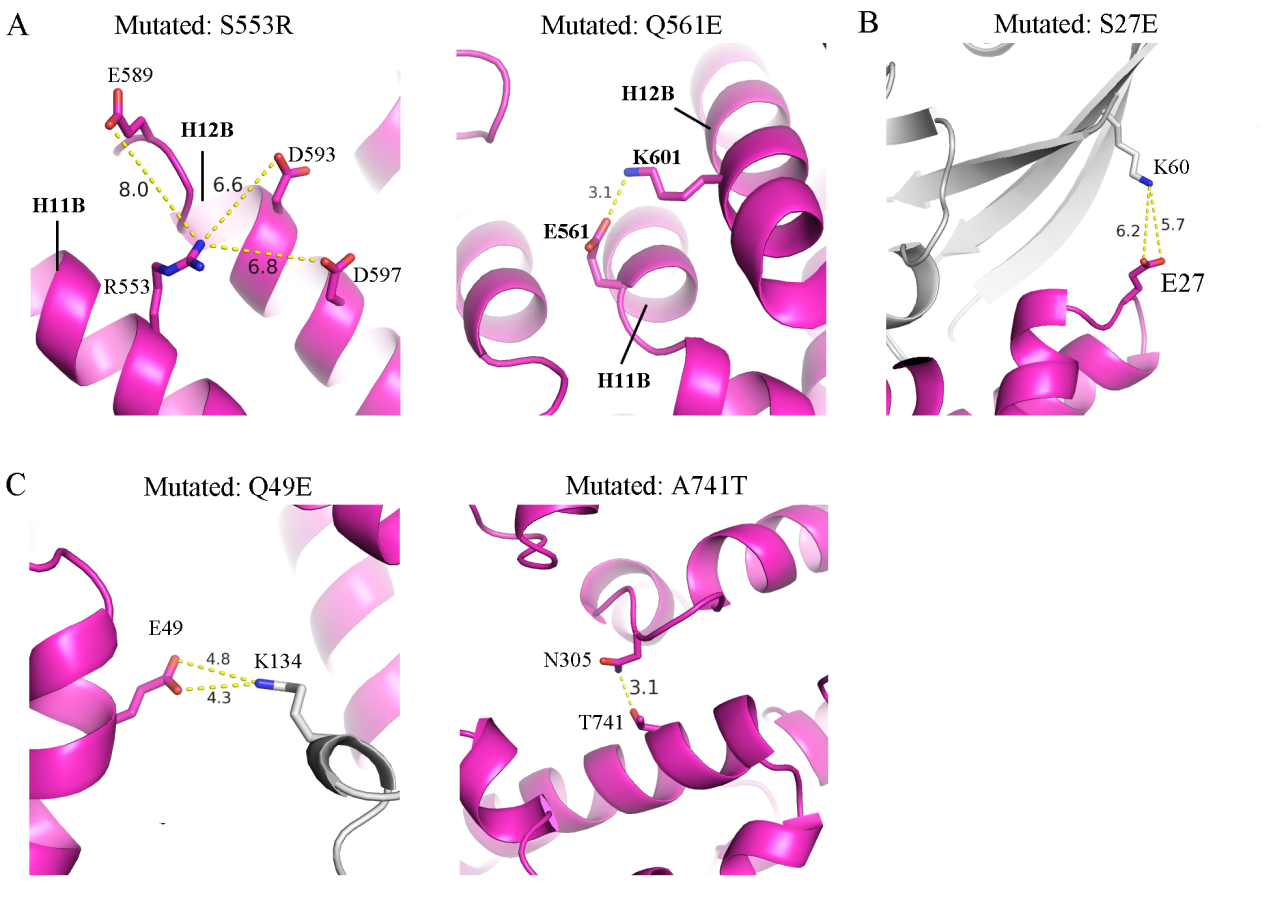

Figure S5. Conformation of mutated residues in the plumbagin-yCRM1e complex. A) residues mutated to open the NES-binding groove. Inter-atomic distances are labelled as dotted lines. B) Residue mutated to improve Ran-CRM1 affinity. C) Residues mutated to improve packing.

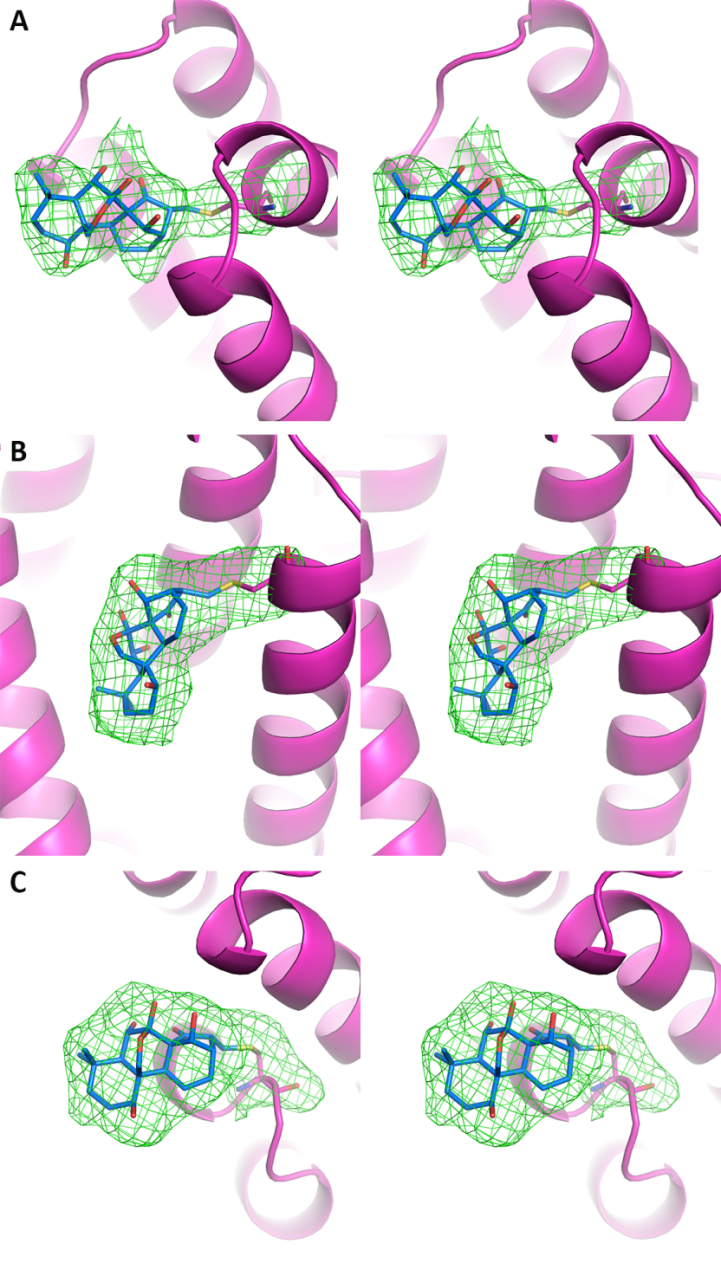

Figure S6. Stereo view of the Fo-Fc SA omit maps (green mesh) contoured at 4σ level. A) omit map for oridonin and the conjugated cysteine C152. CRM1 is shown as magenta cartoon representation. Oridonin is shown as cyan sticks. B) omit map for oridonin and the conjugated C539. C) omit map for oridonin and the conjugated C1022.

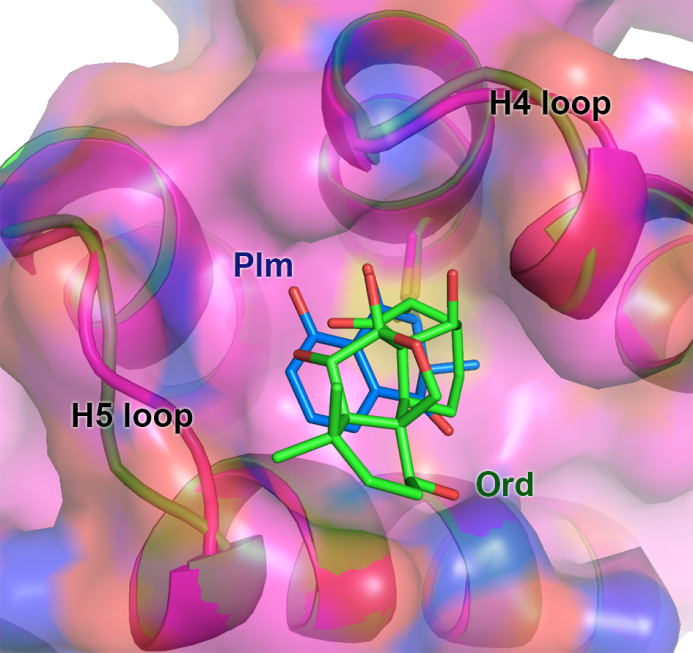

Figure S7. Overlay of C-152 bound oridonin (green) and CRM1a-C152 bound plumbagin (blue). Some differences of H4 and H5 loops are observed.

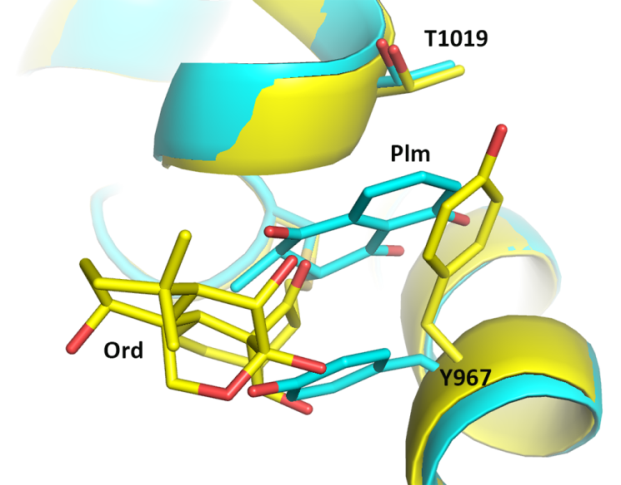

Figure S8. Comparison of C1022-linked plumbagin and oridonin. The CRM1 coloring is different from previous figures to illustrate side chain changes of Y967 and T1019. Oridonin is not inserted into the plumbagin-binding channel formed by Y967 and T1019. In oridonin complex, Y967 shifts and touches T1019, closing the plumbagin-binding channel.

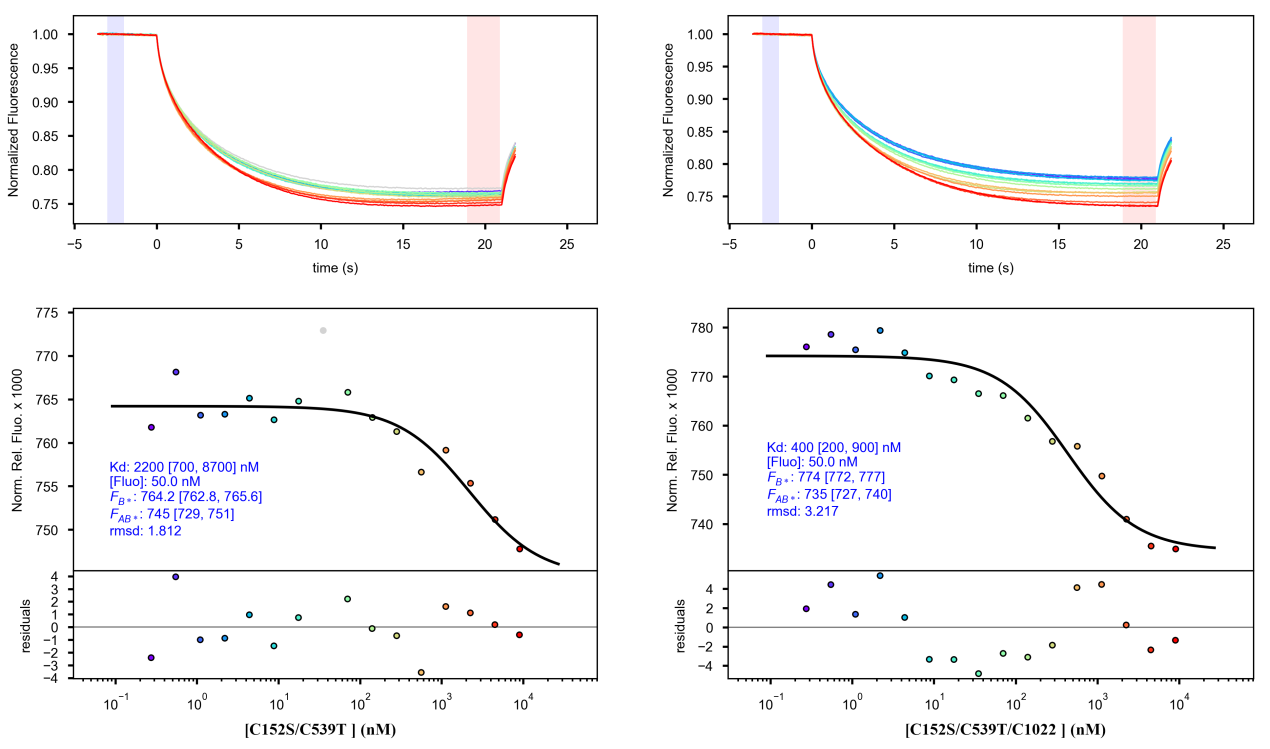

Figure S9. Microscale thermophoresis data of C152S/C539T double mutant or C152S/C539T/C1022S triple mutant binding to MBP-NES^MVM^ in the presence of excess (60 µM) oridonin. The data is processed using the PALMIST programme. MBP-NES^MVM^ was labelled with RED-NHS dye and kept at a constant concentration of 50 nM. The LED power and MST power are 25% and 40% respectively. The assay buffer contains 50 mM Hepes pH 7.5, 200 mM NaCl, 0.05% Tween-20.

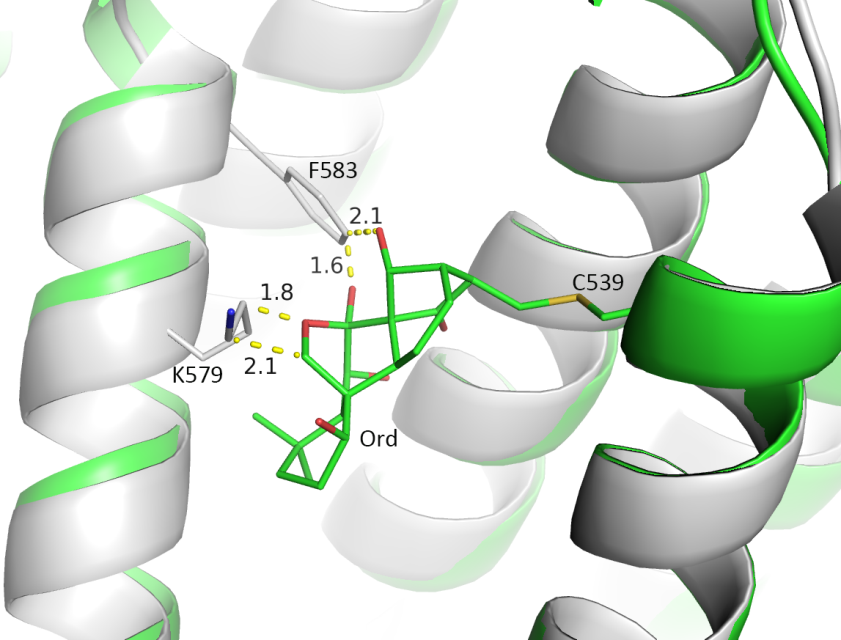

Figure S10. Alignment of NES-bound groove (grey, 6CIT) with CRM1e (green) using the H11A residues (529-540). Oridonin clashes (yellow dash lines) with F583 and K579 on H12A in the NES-bound structure.

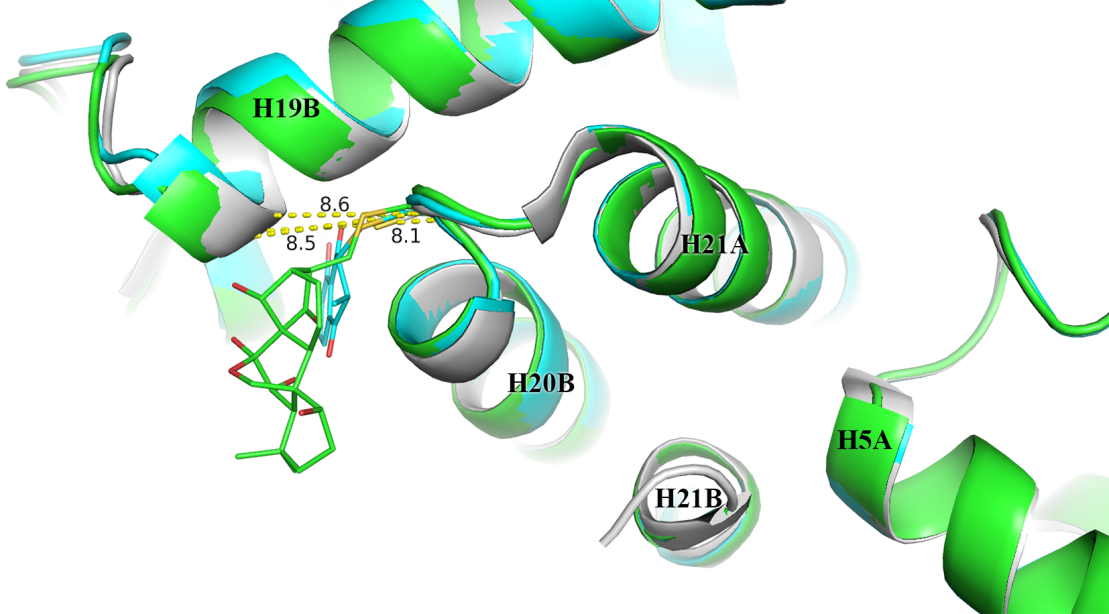

Figure S11. Potential mechanism of NES-inhibition by C1022 conjugation. NES-bound structure (6CIT, grey) is superimposed with plumbagin (cyan) and oridonin (green) structures. C1022 conjugations by plumbagin or oridonin slightly increased the distance between H20B and H19B (8.1Å to 8.5Å or 8.6 Å). The changes could be larger in solution, since crystal packing might have masked a portion of the changes. The C1022-conjugation might potentially disrupt or weaken the interaction between H21 and H5A, which was previously shown to be important for NES binding (see discussion).
